# MsQuality – an interoperable open-source package for the calculation of standardized quality metrics of mass spectrometry data

**DOI:** 10.1101/2023.05.12.540477

**Authors:** Thomas Naake, Johannes Rainer, Wolfgang Huber

## Abstract

**Motivation:** Multiple factors can impact accuracy and reproducibility of mass spectrometry data. There is a need to integrate quality assessment and control into data analytic workflows.

**Results:** The `MsQuality` package calculates 40 low-level quality metrics based on the controlled mzQC vocabulary defined by the HUPO-PSI on a single mass spectrometry-based measurement of a sample. It helps to identify low-quality measurements and track data quality. Its use of community-standard quality metrics facilitates comparability of quality assessment and control (QA/QC) criteria across datasets.

**Availability:** The R package `MsQuality` is available through Bioconductor at https://bioconductor.org/packages/MsQuality.

**Supplementary information:** Supplementary data are available online.

---

Mass spectrometry (MS) is a versatile analytical technique that has been adopted in a variety of disciplines, including proteomics, metabolomics, and lipidomics, enabling the identification and quantification of a wide range of molecules. Obtaining high-quality data from mass spectrometry experiments can be a challenging task, as numerous factors can impact the accuracy and reproducibility of the obtained data. To ensure that MS data is fit for purpose, quality assessment and quality control (QA/QC) need to be performed close to data production from raw data (Köcher *et al*., 2011; Bereman, 2015). Use of standardized quality metrics described by a controlled vocabulary helps in making QA/QC more comparable across datasets and data producers and increases transparency and trustworthiness of such measures as viewed by data users (Mayer *et al*., 2012, 2013).

Here, we introduce the `MsQuality` R-package, which provides functionality to calculate, assess, and track quality metrics for mass spectrometry-derived spectral data of a single massspectrometry-based measurement of a sample. The package provides 40 of the mzQC quality metrics defined by the Human Proteome Organization-Proteomics Standards Initiative (HUPO-PSI, hupo-psi.github.io/mzQC). These are calculated on low-level MS data such as retention times, *m/z*, and associated intensity values. The package automates tracking and quantification of data quality and helps to integrate these computations in routine workflows, thereby, `MsQuality` facilitates the identification of measurements with low quality, including those with a high occurrence of missing values, ahead-of-time termination of chromatographic runs, or low instrument sensitivity.

Following the definitions by Bittremieux *et al*. (2017), `MsQuality` focuses on the calculation of inter-experiment metrics, which is a summarization of an intra-experiment metric. Examples for intra-experiment metrics are the chromatogram of the total ion current (TIC) over the retention time. Inter-experiment metrics, on the other hand, facilitate the comparison of multiple MS runs or experiments, e.g., via longitudinal analysis of quality metrics, such as the fractions of the total retention time required to accumulate a given percentile of the TIC.

## 1 Usage scenario and implementation

`MsQuality` offers easy-to-use means of evaluating data quality on a per-measurement basis, including the identification of low-quality measurements, biases and outliers, variations in calibration, and batch and confounding effects within datasets (Fig. 1 a and b). Its use of community standards for data representation in mass spectrometry defined by HUPO-PSI facilitates comparison, consistent storage, reporting and exchange of quality metrics and quality control criteria.

**Figure 1:**
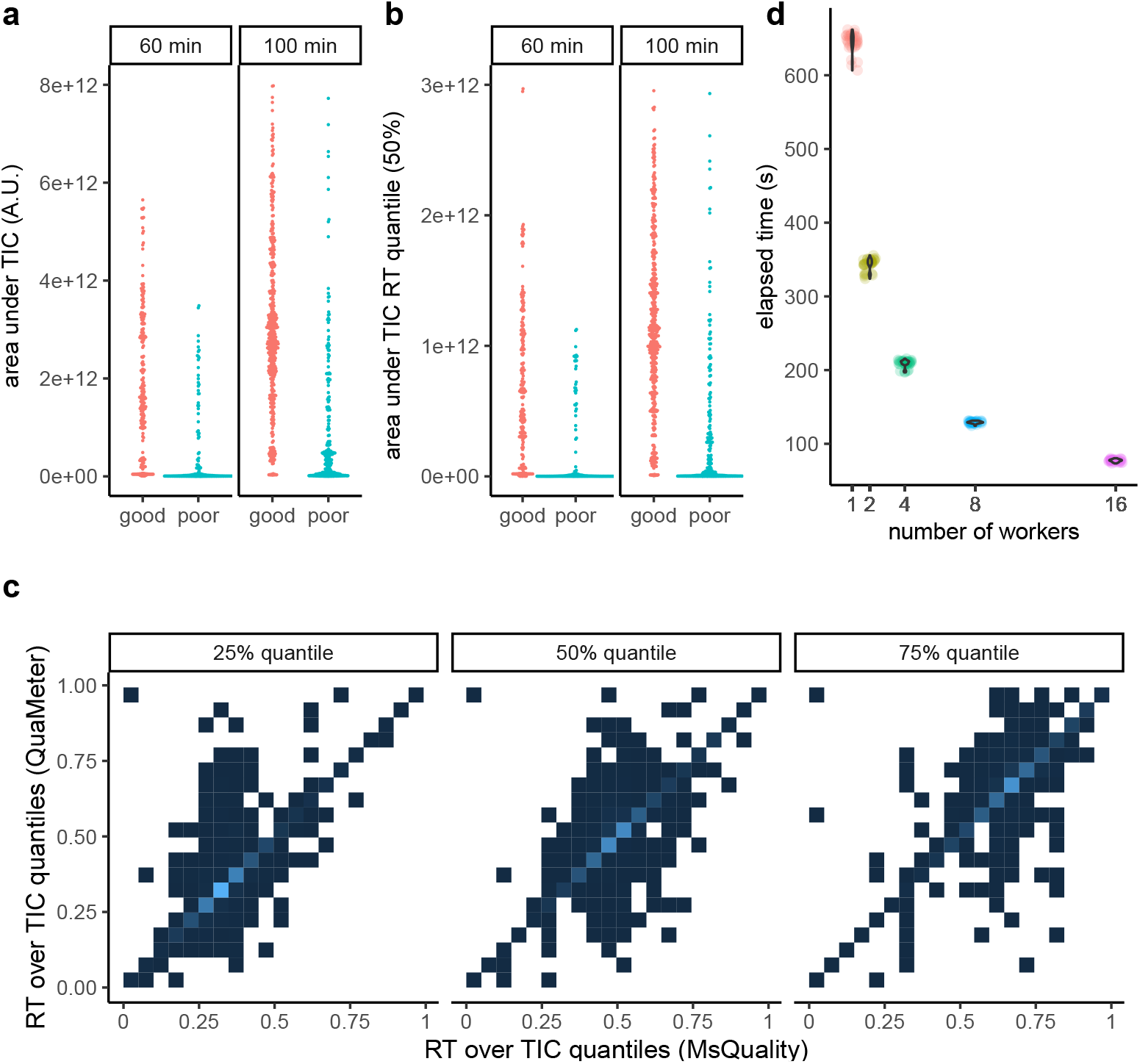
Examples of `MsQuality` functionality. Metrics are based on MS1 spectra; one data point is obtained per MS1 spectrum. (a) Area under TIC: The area under the total ion chromatogram. (b) Quantiles of area under the total ion chromatogram of the retention time (TIC RT), here, the 50% quantile. For (a) and (b) the data points are displayed in a beeswarm plot and stratified for high-quality and low-quality measurements as classified in Amidan *et al*. (2014). (c) Comparison of quality metrics calculated by `MsQuality` and QuaMeter: RT over TIC quantiles. The data points are displayed as 2D densities. Brighter areas corresond to high 2D density areas. (d) Wall-clock execution time for the calculation of quality metrics of the data set of Amidan *et al*. (2014) when parallel computing is used (1, 2, 4, 8, and 16 workers). A.U. arbitrary units.

The versatility of `MsQuality` in calculating metrics extends to a wide range of applications, from small-scale studies to long-term acquisition of mass spectrometry data, e.g. a core facility running an instrument for months and years. We demonstrate the utility of `MsQuality` in two case studies: a dataset of 180 cancer cell lines obtained by flow injection analysis (Cherkaoui *et al*., 2022) and a liquid chromatography (LC)-MS dataset of the same control sample (Amidan *et al*., 2014) as instance of a long-term quality control usage scenario. The values computed by `MsQuality` agree with those of QuaMeter (Ma *et al*., 2012) (Fig. 1 c): 75% of the analyzed `MsQuality` metrics showed Pearson correlation coefficients over 0.81 and Spearman correlation coefficients over 0.87 (see the Supplementary Data for further details).

`MsQuality` is implemented as an GPL-3-licensed open-source R package, building upon the established `Spectra` and `MsExperiment` packages (Rainer *et al*., 2022) to provide and represent the MS data. Thus, `MsQuality` supports a large variety of data input formats as well as analyses of very large experiments through the use of data representations with low memory footprint. Native parallelization enables a fast and scalable calculation of quality metrics (Fig. 1 d, see the Supplementary Data for further details).

Finally, `MsQuality` requires little programmatic interaction and is designed to be userfriendly. After the instantiation of `Spectra` or `MsExperiment` object, a single function call is needed to calculate the quality metrics.

## 2 Conclusion

The `MsQuality` R-package provides functionality to calculate, assess, and track quality metrics for mass spectrometry-derived spectral data. It offers easy-to-use means of evaluating data quality on a per-measurement basis, enabling researchers the identification of low-quality measurements. By using standardized quality metrics via the controlled vocabulary of HUPO-PSI, `MsQuality` helps to make QA/QC more comparable across datasets and data producers. The implementation of `MsQuality`’s metric calculation is designed to be user-friendly and streamlined and requires little programmatic interaction, facilitating reproducible calculation and evaluation of data quality metrics. `MsQuality` contributes to the expanding list of tools that use the `Spectra/MsExperiment` framework (Rainer *et al*., 2022) to address various stages in the analysis pipeline of mass spectrometry data. By building upon this extensive ecosystem for mass spectrometry data, `MsQuality` enables researchers to create seamless analysis workflows for rapid, efficient, and standardized evaluation of MS data quality, ultimately leading to more robust scientific discoveries in mass spectrometry workflows.

## 3 Acknowledgements

We acknowledge feedback from Friedemann Ringwald, Hagen Gegner, and Torsten Müller on usability of `MsQuality` and all developers and maintainers of the R/Bioconductor packages `MsQuality` is built upon. We would like to thank Nicola Zamboni for his valuable assistance in locating and understanding the data of Cherkaoui *et al*. (2022).

## 3.1 Author contributions statement

T.N. conceptualized the project. T.N. and J.R. implemented the algorithms as an R package.

T.N. analysed the results. W.H. provided feedback and guidance. T.N., J.R. and W.H. wrote the manuscript.

