## Supplementary File for "MsQuality – an interoperable open-source package for the calculation of standardized quality metrics of mass spectrometry data"

Supplementary Data

Thomas Naake* Johannes Rainer† Wolfgang Huber‡

**4 Session info** **24**

*Genome Biology Unit, European Molecular Biology Laboratory, Meyerhofstrasse 1, 69117 Heidelberg, Germany

†Institute for Biomedicine (Affiliated Institute of the University of Lübeck), Eurac Research, Viale Druso 1, 39100 Bolzano, Italy

‡Genome Biology Unit, European Molecular Biology Laboratory, Meyerhofstrasse 1, 69117 Heidelberg, Germany

This is the Supplementary Information for the publication "MsQuality - an interoperable open-source package for the calculation of standardized quality metrics of mass spectrometry data". It demonstrates
functionality of the `MsQuality` package on two example analysis workflows using the data sets of Cherkaoui et al. [2022] and Amidan et al. [2014].

The `MsQuality` package calculates low-level quality metrics that only require minimal information about the mass spectrometry data: retention time, m/z values, and associated intensities. The list of quality metrics provided by the mzQC framework (hupo-psi.github.io/mzQC) is extensive, also including metrics that depend on
higher level information which might not be readily accessible from .raw or .mzML files, such as pump pressure mean, or that rely on alignment results, like retention time mean shift, signal-to-noise ratio, precursor errors (ppm). Such metrics are currently not implemented in `MsQuality`.

The `MsQuality` package relies on the `Spectra` package for data import and representation. Quality metrics are calculated from the information in a `Spectra` object. The `dataOrigin` variable is used to distinguish between the MS data from different measurements/files. Section 1 loads these and other packages into the environment of the `R` session in order to run all analyses.

In subsequent sections of this document the quality of two data sets will be analyzed:

- Section 2: the Cherkaoui et al. [2022] data set is a mass spectrometry (MS) metabolomics data set of 180 cancer cell lines obtained via flow injection analysis (TOF, negative ionization mode). The data set comprises a total of 1397 measurements.
- In Section 3: the Amidan et al. [2014] data set consists of 3431 LC-MS measurements of a single QC sample (whole cell lysate of *Shewanella oneidensis*). The QC sample was measured on Exactive, LTQ IonTrap, LTQ Orbitrap, and Velos Orbitrap instruments.

We note that these quality metrics are indicative, but by themselves might not be sufficient for data quality control decision-making, such as removing low-quality measurements, which might require additional consideration of more advanced analytics, such as those provided by `MatrixQCvis` [Naake and Huber, 2022]. As stated previously [Bittremieux et al., 2017], the utility of QC metrics depend on the type of the sample, e.g., on whether a single peptide or a complex lysate of proteins is analyzed [Bereman, 2015, Köcher et al. [2011],Paulovich et al. [2010]].

In this document, we will - create `Spectra` objects from the raw data of the two datasets, - calculate the quality metrics on these data sets, - visualize some of the metrics, and - assess performance and scalability of the implemented algorithms using the `microbenchmark` package.

Due to journal's publication format, this document presents static plots. Note that the `MsQuality` package also includes an interactive shiny application to interactively navigate quality metrics, with plots based on the plotly framework. For reproducibility, we provide the source .Rmd file in the accompanying GitHub repository.

A list of the attached pacakges can be found in Section 4. We will indicate which parts of this document are reproducible.

### 1 Preparation of the environment

This analysis uses functions from multiple `R` packages, including `Spectra` for representing mass spectrometry spectral data and `MsQuality` for calculating quality metrics. Other packages are required for data visualization (`ggplot2`, `ggbeeswarm`, `ggpubr`), data wrangling (`dplyr`, `readxl`, `stringr`, `tibble`, `tidyr`), and performance and scalability analysis (`microbenchmark`). Before starting the analysis, ensure to load these packages.

```
## load packages for visualization
library("ggplot2")
library("ggbeeswarm")
library("ggpubr")

## load packages for data wrangling
library("dplyr")
library("readxl")
library("stringr")
library("tibble")
library("tidyr")

## load packages for performance and scalability analysis
library("microbenchmark")

## load packages for storing spectral data and calculating quality metrics
library("Spectra")
library("MsQuality")
```

### 2 Cherkaoui et al. [2022]: A functional analysis of 180 cancer cell lines reveals conserved intrinsic metabolic programs

The .mzML files were downloaded from the MassIVE database (accession number MSV000087155, available at https://massive.ucsd.edu/) via ftp://massive.ucsd.edu/MSV000087155.

We use the BiocFileCache package from Bioconductor to download and cache the mzML files locally. To this end we first determine below the full file names of all mzML files of this data set.

All parts in the section *Cherkaoui et al. [2022]: A functional analysis of 180 cancer cell lines reveals conserved intrinsic metabolic programs* are reproducible except the parallelization steps in the subsection **Performance under parallelization**. For these steps precalculated objects are loaded to the environment.

```
url <- "ftp://massive.ucsd.edu/MSV000087155/ccms_peak/New_mzMLFinal/"
library(curl)
```

```
## Using libcurl 7.84.0 with Schannel
```

```
file_list <- readLines(curl(url, "r"))
ftp_files <- strsplit(file_list, " +")
ftp_files <- vapply(
    ftp_files, function(z) paste0(tail(z, 2), collapse = " "), character(1))
```

With the file names available we next create a *BiocFileCache* instance adding the files. The *BiocFileCache* will take care of downloading files that are not already available in the local cache thus preventing unneeded downloads.

```
library(BiocFileCache)
## every additional result should be saved in there
cherkaoui <- BiocFileCache("../Cherkaoui2022", ask = FALSE)
path <- bfcrpath(cherkaoui, paste0(url, curl_escape(ftp_files)))
```

These downloaded mzML files can however not be directly loaded because they are not fully compliant with the open mzML standard file format (internal references to instrumentation configuration are missing). We thus need to process all files to remove these incompatible lines from each mzML file. This needs to be done (once) using the below unix shell commands that should be executed in the folder containing the downloaded files.

```
cd ../Cherkaoui2022
sed -i 's/<run defaultInstrumentConfigurationRef=.*/<run/g' *.mzML
sed -i '/^<scanWindowList/,/^<\/scanWindowList/d' *.mzML
```

#### 2.1 Instantiation of the Spectra object

In the subsequent analysis, a `Spectra` object is instantiated. The operations were executed within a (high-performance) computing environment (31 cores, 64 GB RAM pool for all cores).

```
## create the Spectra object
sps <- Spectra(path, backend = MsBackendMzR())
```

#### 2.2 Calculate the metrics via `MsQuality`

`MsQuality` uses the `Spectra` class for storing the spectral data. In this particular case, where the spectral data was obtained via flow injection analysis, metrics that incorporate retention time information are not relevant and the analysis will only focus on the three metrics

- `numberSpectra`, **Number of MS1 spectra** (QC:4000059), "The number of MS1 events in the run." [PSI:QC];
- `areaUnderTic`, **Area under TIC** (QC:4000077), "The area under the total ion chromatogram." [PSI:QC];
- `mzAcquisitionRange`, **m/z acquisition range** (QC:4000138), "Upper and lower limit of m/z values at which spectra are recorded." [PSI:QC].

The metrics are calculated using the function `calculateMetricsFromSpectra`, which takes as input the `Spectra` object, `sps`, and the above-defined metrics. Optional parameters can also be passed to this function for further control of the calculation, such as `msLevel` for cases where multiple mass spectra levels are present in the `Spectra` object. It is unnecessary to specify `msLevel` in the current context since only MS1 level spectra are stored in the `Spectra` object.

```
metrics <- c("numberSpectra", "areaUnderTic", "mzAcquisitionRange")

metrics_sps <- calculateMetricsFromSpectra(spectra = sps,
    metrics = metrics)
```

#### 2.3 Visualization

We next visualize the three quality metrics using the `ggplot2` package. We include also information from the original study Cherkaoui et al. [2022] in particular which of the files were included in the final analysis. The results of the study are available from this resource: https://doi.org/10.3929/ethz-b-000511784 . We first download and cache the *PrimaryAnalysis.zip* archive that contains all results, unzip it to a temporary folder and import the *metabolomics_180CCL.xlsx* file.

```
l <- paste0("https://www.research-collection.ethz.ch/bitstream/handle/",
            "20.500.11850/511784/PrimaryAnalysis.zip?sequence=1&isAllowed=y")
arch <- bfcrpath(cherkaoui, l)
unzip(zipfile = arch, files = "PrimaryAnalysis/Metabolomics_180CCL.xlsx",
```

```
      exdir = tempdir())
measurements <- read_xlsx(
    file.path(tempdir(), "PrimaryAnalysis/Metabolomics_180CCL.xlsx"),
    sheet = "injections")
```

From this excel sheet we retrieve the information whether a measurements was analyzed or ecluded and add this information to the `metrics_sps` object with the quality information.

We then create a Figure to compare the differences in quality metrics between the analyzed and excluded measurements (Figure S1).

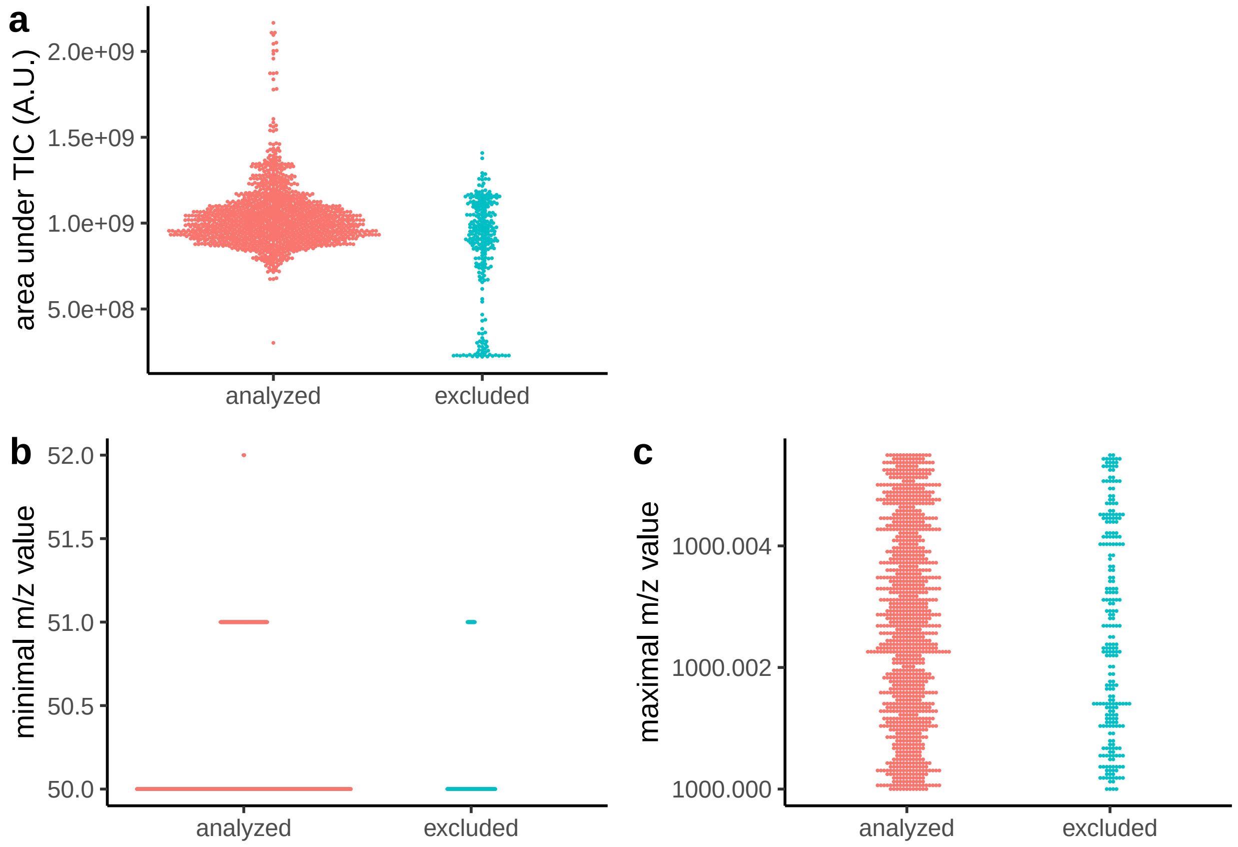

Figure S1: Quality metrics for data set of Cherkaoui et al. [2022] stratified by information if the measurement was analyzed (`yes`) or excluded (`no`). (a) Area under the TIC (`areaUnderTic`). (b) Minimum values of the m/z acquisition range (`mzAcquisitionRange.min`). (c) maximum values of the m/z acquisition range `mzAcquisitionRange.max`). A.U.: arbitrary units.

Figure S1 demonstrates that the excluded measurements show a bimodal distribution of the total ion current (TIC). Specifically, some of the excluded measurements have lower total ion current (TIC) values, which was already noted in the original publication and was the reason for their exclusion from subsequent analysis steps. Figure S1 a serves as a visual confirmation of this statement and aids in understanding the data quality of the measurements The metrics `mzAcquisitionRange.max` and `mzAcquisitionRange.min` on the other hand (Figure S1 (b) and (c)) are not informative for the decision making on excluding/including the measurements in further analysis steps.

#### 2.4 Performance under parallelization

An important aspect, especially when dealing with large amount of data, is scalability and performance when computing the quality metric.

By monitoring parallelization, it is possible to determine the scalability of the computation and ensure that the performance of the analysis remains acceptable as the data size increases.

We measure below the time it takes to evaluate the calculation of quality metrics by parallelizing the tasks on 1, 2, 4, 8, and 16 workers using the `microbenchmark` package. This package allows for precise measurement of the execution time of `R` expressions by repeating the evaluation multiple times and providing detailed summary statistics of the execution times.

```
metrics <- c("numberSpectra", "areaUnderTic", "mzAcquisitionRange")
df_mb <- microbenchmark(calculateMetricsFromSpectra(spectra = sps,
        metrics = metrics, BPPARAM = MulticoreParam(workers = 1)),
    workers_2 = calculateMetricsFromSpectra(spectra = sps,
        metrics = metrics, BPPARAM = MulticoreParam(workers = 2)),
    workers_4 = calculateMetricsFromSpectra(spectra = sps,
        metrics = metrics, BPPARAM = MulticoreParam(workers = 4)),
    workers_8 = calculateMetricsFromSpectra(spectra = sps,
        metrics = metrics, BPPARAM = MulticoreParam(workers = 8)),
    workers_16 = calculateMetricsFromSpectra(spectra = sps,
        metrics = metrics, BPPARAM = MulticoreParam(workers = 16)),
    times = 110L, control = list(warmup = 10), check = "equal"
)
```

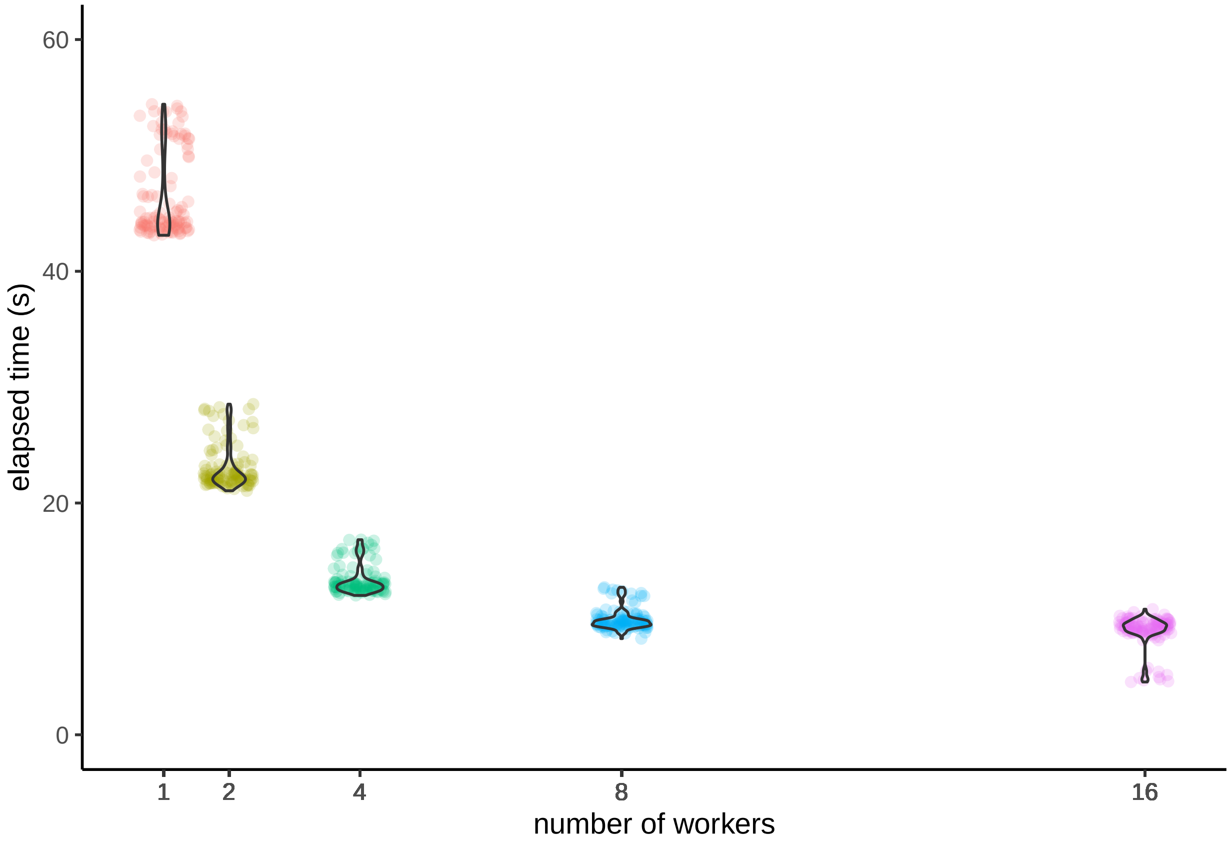

Figure S2: Execution time for the calculation of quality metrics of the data set of Cherkaoui et al. [2022] under parallelization (1, 2, 4, 8, and 16 workers).

By parallelizing the calculation of the quality metrics across multiple workers, it is possible to significantly reduce the execution time, and the `microbenchmark` package was used to accurately measure the performance improvements achieved by parallelization (Figure S2). The parallelization process can help in the management of bigger data sets, and to save valuable time in data analysis.

### 3 Amidan et al. [2014]: Signatures for mass spectrometry data quality

The RAW files were downloaded from

- ftp://ftp.pride.ebi.ac.uk/pride/data/archive/2013/10/PXD000320 (`1_of_5`),
- ftp://ftp.pride.ebi.ac.uk/pride/data/archive/2013/10/PXD000321 (`2_of_5`),
- ftp://ftp.pride.ebi.ac.uk/pride/data/archive/2013/10/PXD000322 (`3_of_5`),
- ftp://ftp.pride.ebi.ac.uk/pride/data/archive/2013/10/PXD000323 (`4_of_5`),
- ftp://ftp.pride.ebi.ac.uk/pride/data/archive/2013/10/PXD000324 (`5_of_5`).

Subsequently, the RAW files were converted into `.mzML` files using MSConvertGUI (64-bit, v3.0.22015-aadd392) with setting `peakPicking` to `vendor msLevel=1-`.

The creation of the Figures in the section *Amidan et al. [2014]: Signatures for mass spectrometry data quality* are reproducible. Due to long computation time or requirement of an environment that enables for parallelization, the creation of the `Spectra` object in the subsection **Instantiation of the Spectra object**, the calculation of the quality metrics in the subsection **Calculate the metrics via `MsQuality`**, and the parallelization steps in the subsection **Performance under parallelization** are precomputed.

#### 3.1 Instantiation of the Spectra object

In the subsequent analysis, a `Spectra` object is instantiated. The operations were executed within a (high-performance) computing environment (31 cores, 128 GB RAM pool for all cores), where the .mzML files were stored in the directory `Amidan2014`.

```
## read the file with protein intensities
path <- "/scratch/naake/Amidan2014"
fls <- dir(path, full.names = TRUE, recursive = TRUE, pattern = "mzML")

## create the Spectra object
sps <- Spectra(fls, backend = MsBackendMzR())
```

#### 3.2 Calculate the metrics via `MsQuality`

`MsQuality` utilizes `Spectra` objects that store the spectral data. Here, retention time information was available from the .mzML files and a higher number of metrics could be calculated:

- *rtDuration*, **RT duration** (QC:4000053), "The retention time duration of the MS run in seconds, similar to the highest scan time minus the lowest scan time." [PSI:QC];
- *rtOverTicQuantiles*, **RT over TIC quantile** (QC:4000054), "The interval when the respective quantile of the TIC accumulates divided by retention time duration. The number of quantiles observed is given by the size of the tuple." [PSI:QC];
- *rtOverMsQuarters*, **MS1 quantiles RT fraction** (QC:4000055), "The interval used for acquisition of the first, second, third, and fourth quarter of all MS1 events divided

by RT-Duration." [PSI:QC];

- *rtOverMsQuarters*, **MS2 quantiles RT fraction** (QC:4000055), "The interval used for acquisition of the first, second, third, and fourth quarter of all MS2 events divided by RT-Duration." [PSI:QC];
- *ticQuartileToQuartileLogRatio*, **MS1 TIC-change quartile ratios** (MS:4000057), "The log ratios of successive TIC-change quartiles. The TIC changes are the list of MS1 total ion current (TIC) value changes from one to the next scan, produced when each MS1 TIC is subtracted from the preceding MS1 TIC. The metric's value triplet represents the log ratio of the TIC-change Q2 to Q1, Q3 to Q2, TIC-change-max to Q3" [PSI:MS], `mode = "TIC_change"`;
- *ticQuartileToQuartileLogRatio*, **MS1 TIC quartile ratios** (MS:4000058), The log ratios of successive TIC quartiles. The metric's value triplet represents the log ratios of TIC-Q2 to TIC-Q1, TIC-Q3 to TIC-Q2, TIC-max to TIC-Q3." [PSI:MS], 'mode = "TIC";
- *numberSpectra*, **Number of MS1 spectra** (QC:4000059), "The number of MS1 events in the run." [PSI:QC];
- *numberSpectra*, **Number of MS2 spectra** (QC:4000060), "The number of MS2 events in the run." [PSI:QC];
- *medianPrecursorMz*, **Precursor median m/z for IDs** (QC:4000065), "Median m/z value for all identified peptides (unique ions) after FDR." [PSI:QC];
- *rtIqr*, **Interquartile RT period for peptide identifications** (QC:4000072), "The interquartile retention time period, in seconds, for all peptide identifications over the complete run." [PSI:QC];
- *rtIqrRate*, **Peptide identification rate of the interquartile RT period** (QC:4000073), "The identification rate of peptides for the interquartile retention time period, in peptides per second." [PSI:QC];
- *areaUnderTic*, **Area under TIC** (QC:4000077), "The area under the total ion chromatogram." [PSI:QC];
- *areaUnderTicRtQuantiles*, **Area under TIC RT quantiles** (QC:4000078), "The area under the total ion chromatogram of the retention time quantiles. Number of quantiles are given by the n-tuple." [PSI:QC];
- *medianTicRtIqr*, **Median of TIC values in the RT range in which the middle half of peptides are identified** (QC:4000130), "Median of TIC values in the RT range in which half of peptides are identified (RT values of Q1 to Q3 of identifications)" [PSI:QC];
- *medianTicOfRtRange*, **Median of TIC values in the shortest RT range in which half of the peptides are identified** (QC:4000132), "Median of TIC values in the shortest RT range in which half of the peptides are identified" [PSI:QC]
- *mzAcquisitionRange*, **m/z acquisition range** (QC:4000138), "Upper and lower limit of m/z values at which spectra are recorded." [PSI:QC];
- *rtAcquisitionRange*, **Retention time acquisition range** (QC:4000139), "Upper and lower limit of time at which spectra are recorded." [PSI:QC];
- *precursorIntensityRange*, **Precursor intensity range** (QC:4000144), "Minimum and maximum precursor intensity recorded." [PSI:QC];
- *precursorIntensityQuartiles*, **Precursor intensity distribution Q1, Q2, Q3**

(QC:4000167), "From the distribution of precursor intensities, the quartiles Q1, Q2, Q3" [PSI:QC];

- *precursorIntensityMean*, **Precursor intensity distribution mean** (QC:4000168), "From the distribution of precursor intensities, the mean." [PSI:QC];
- *precursorIntensitySd*, **Precursor intensity distribution sigma** (QC:4000169), "From the distribution of precursor intensities, the sigma value." [PSI:QC];
- *msSignal10xChange*, **MS1 signal jump (10x) count** (QC:4000172), "The count of MS1 signal jump (spectra sum) by a factor of ten or more (10x) between two subsequent scans" [PSI:QC];
- *RatioCharge1over2*, **Charged peptides ratio 1+ over 2+** (QC:4000174), "Ratio of 1+ peptide count over 2+ peptide count in identified spectra" [PSI:QC];
- *RatioCharge1over2*, **Charged spectra ratio 1+ over 2+** (QC:4000179), "Ratio of 1+ spectra count over 2+ spectra count in all MS2" [PSI:QC];
- *RatioCharge3over2*, **Charged peptides ratio 3+ over 2+** (QC:4000175), "Ratio of 3+ peptide count over 2+ peptide count in identified spectra" [PSI:QC];
- *RatioCharge3over2*, **Charged spectra ratio 3+ over 2+** (QC:4000180), "Ratio of 3+ peptide count over 2+ peptide count in all MS2" [PSI:QC];
- *RatioCharge4over2*, **Charged peptides ratio 4+ over 2+** (QC:4000176), "Ratio of 4+ peptide count over 2+ peptide count in identified spectra" [PSI:QC];
- *RatioCharge4over2*, **Charged spectra ratio 4+ over 2+** (QC:4000181), "Ratio of 4+ peptide count over 2+ peptide count in all MS2" [PSI:QC];
- *meanCharge*, **Mean charge in identified spectra** (QC:4000177), "Mean charge in identified spectra" [PSI:QC];
- *meanCharge*, **Mean precursor charge in all MS2** (QC:4000182), "Mean precursor charge in all MS2" [PSI:QC];
- *medianCharge*, **Median charge in identified spectra** (QC:4000178), "Median charge in identified spectra" [PSI:QC];
- *medianCharge*, **Median precursor charge in all MS2** (QC:4000183), "Median precursor charge in all MS2" [PSI:QC].

The metrics are calculated using the function `calculateMetricsFromSpectra`, which takes as input the `Spectra` object, `sps`, and the above-defined metrics. We calculate the metrics separately for the MS1 (`msLevel 1`) and MS2 spectra (`msLevel 2`).

```
metrics <- c("rtDuration", "rtOverTicQuantiles", "rtOverMsQuarters",
    "ticQuartileToQuartileLogRatio", "numberSpectra",
    "medianPrecursorMz", "rtIqr", "rtIqrRate", "areaUnderTic",
    "areaUnderTicRtQuantiles", "medianTicRtIqr", "medianTicOfRtRange",
    "mzAcquisitionRange", "rtAcquisitionRange", "precursorIntensityRange",
    "precursorIntensityQuartiles", "precursorIntensityMean",
    "precursorIntensitySd", "msSignal10xChange", "ratioCharge1over2",
    "ratioCharge3over2", "ratioCharge4over2", "meanCharge",
    "medianCharge")

metrics_sps_msLevel1 <- calculateMetricsFromSpectra(spectra = sps,
```

```
    metrics = metrics, msLevel = 1L)
metrics_sps_msLevel1 <- calculateMetricsFromSpectra(spectra = sps,
    metrics = metrics, msLevel = 2L)
```

Overall, this function provides a flexible and efficient way to analyze large amounts of mass spectrometry data and obtain insights on the quality of the data.

#### 3.3 Visualization

In the analysis of the Amidan et al. [2014] study, the quality metrics were visualized using the `ggplot2` package. The XLS files pr401143e_si_002.xls and pr401143e_si_003.xls (provided as Supplemental Material of the original publication) was used to extract information on the measurement quality. This information was added to the `metrics_sps_msLevel1` and `metrics_sps_msLevel2` objects.

The Figures S3, S4, and S5 were created as examples to compare the differences between the low- and high-quality measurements for several of the supported quality metrics.

While the metric `rtDuration` (retention time) is a continuous variable, for visualization purposes we will bin the variable to discrete values and will use the measurements over 60 min and 100 min for visualization.

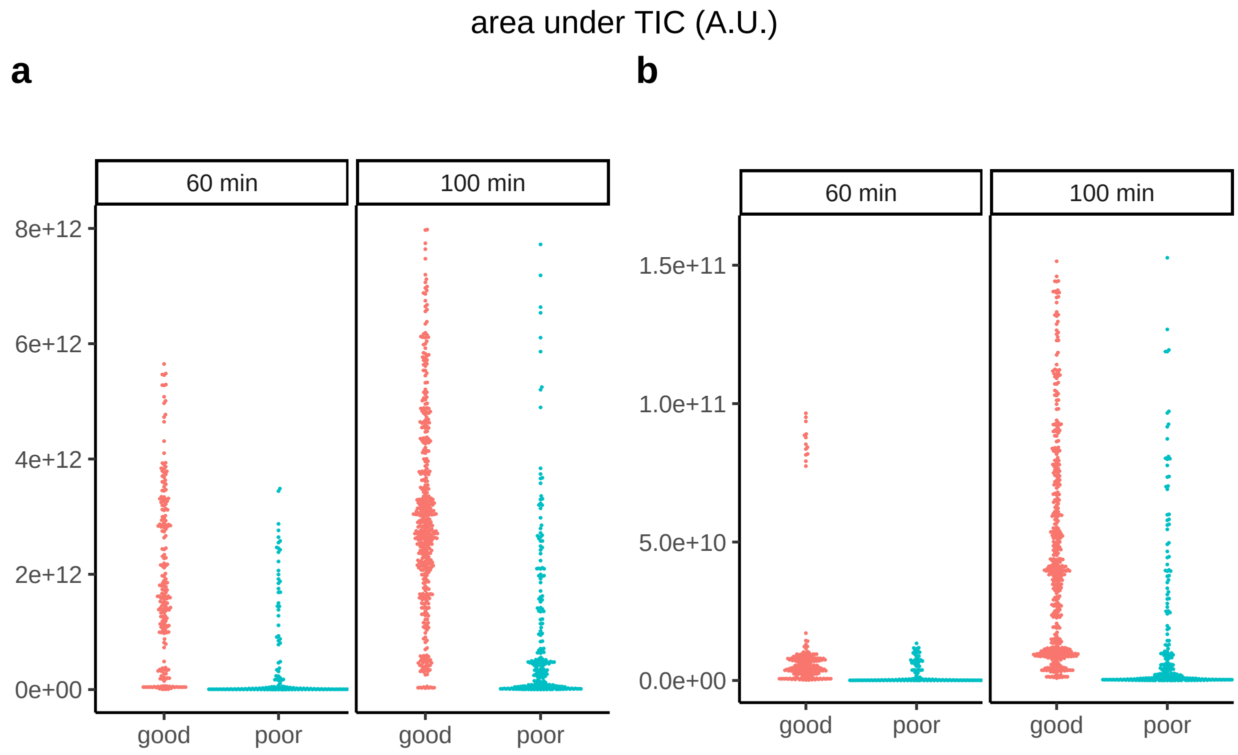

Figure S3: Quality metrics by `MsQuality`: Area under TIC (`areaUnderTic`). The `MsQuality` metrics are calculated from MS1 and MS2 spectra. One data point is obtained per MS1 and MS2 measurement run and the data points are displayed as beeswarm plots stratified for high-quality and low-quality measurements as classified in Amidan et al. [2014]. (a) Area under TIC for MS1 spectra. (b) Area under TIC for MS2 spectra. A.U.: arbitrary units.

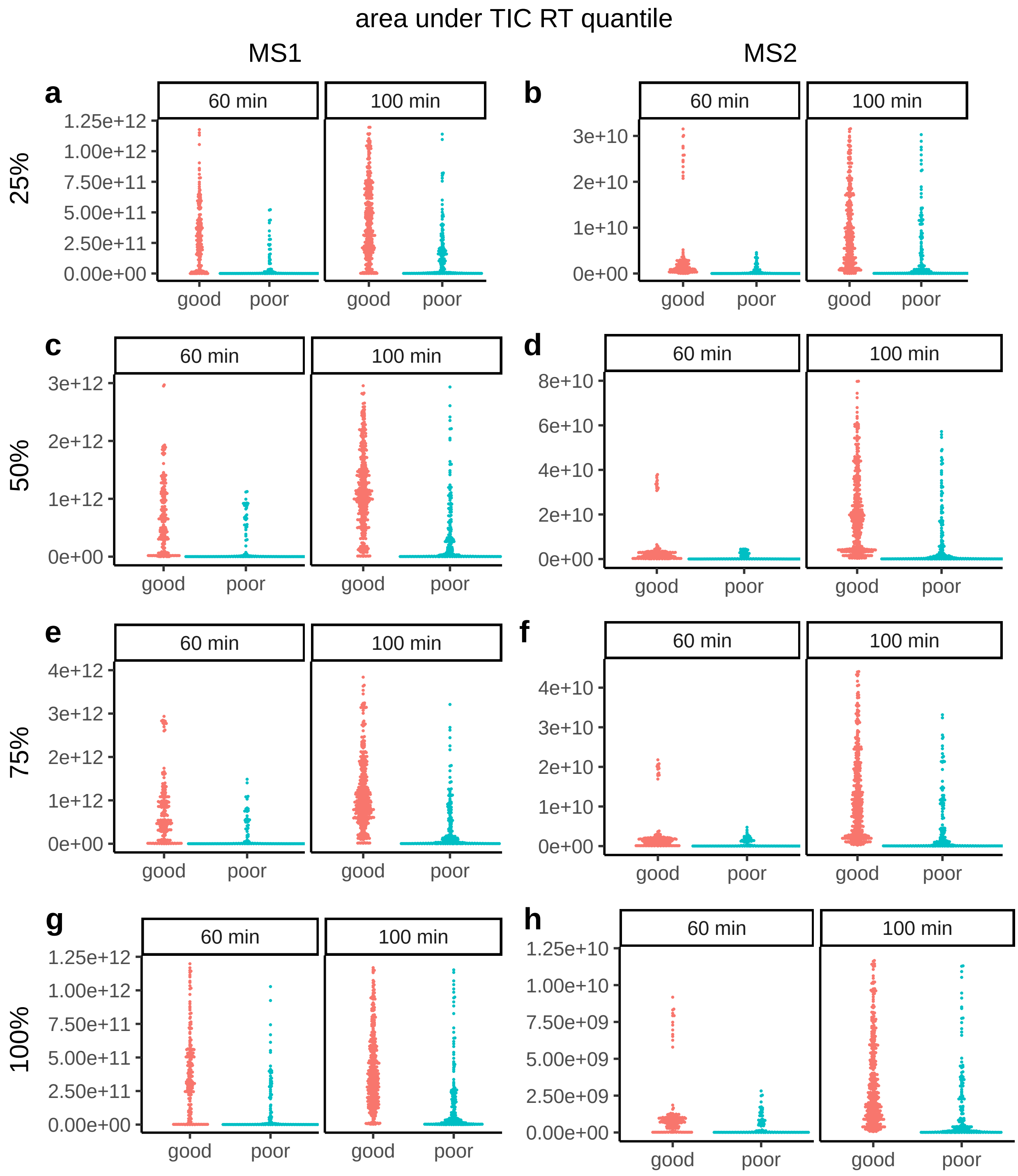

Figure S4: Quality metrics by `MsQuality`: Area under TIC RT quantiles (`areaUnderTicRtQuantiles`). The `MsQuality` metrics are calculated from MS1 and MS2 spectra. One data point is obtained per MS1 and MS2 measurement run and the data points are displayed as beeswarm plots stratified for high-quality and low-quality measurements as classified in Amidan et al. [2014]. (a) 25% quantile for MS1 spectra. (b) 25% quantile for MS2 spectra. (c) 50% quantile for MS1 spectra. (d) 50% quantile for MS2 spectra.E: 75% quantile for MS1 spectra. (f) 75% quantile for MS2 spectra. (g) 100% quantile for MS1 spectra. (h) 100% quantile for MS2 spectra.

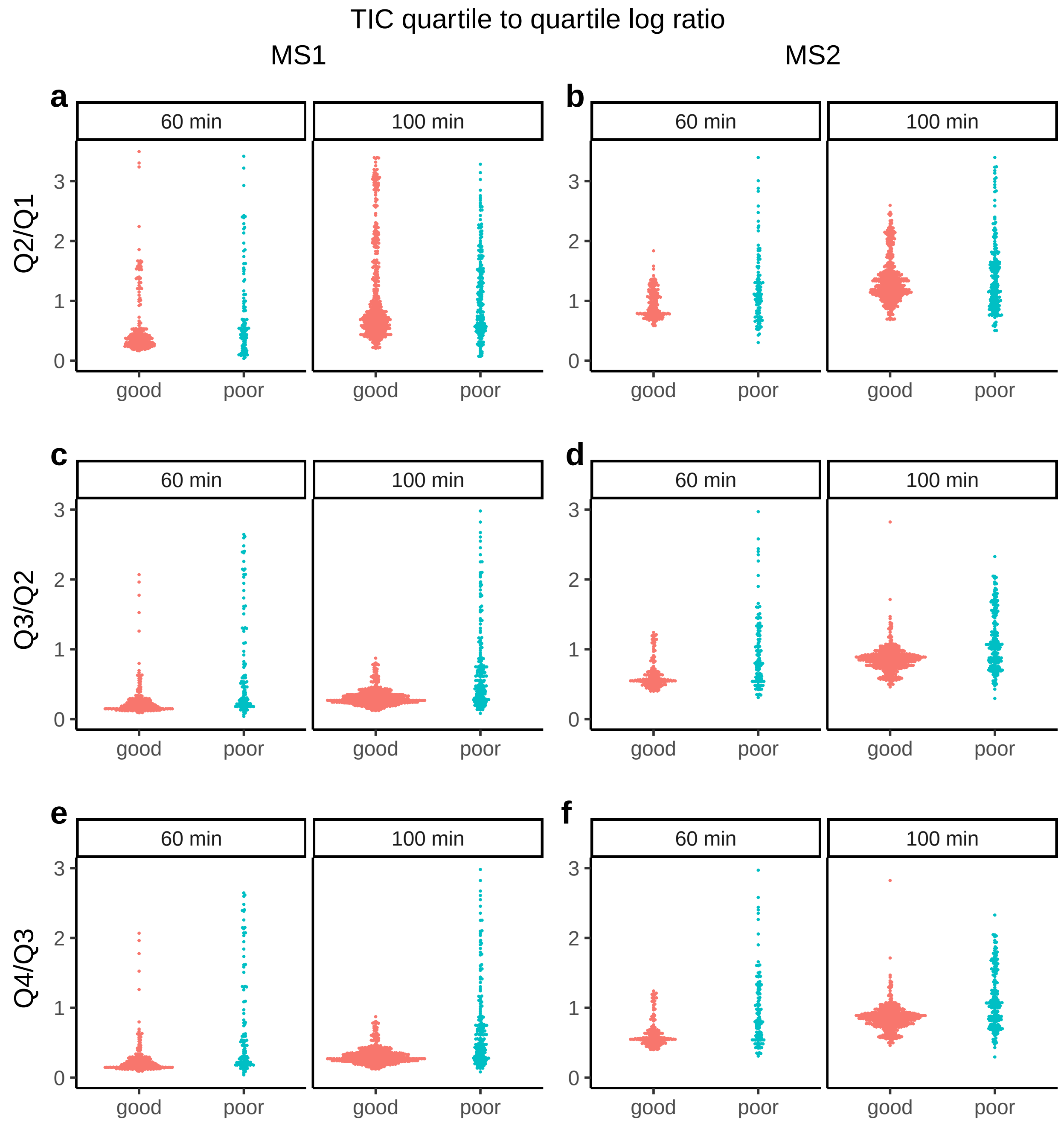

Figure S5: Quality metrics by `MsQuality`: TIC quartile to quartile log ratio (`ticQuartileToQuartileLogRatio`). The `MsQuality` metrics are calculated from MS1 and MS2 spectra. One data point is obtained per MS1 and MS2 measurement run and the data points are displayed as beeswarm plots stratified for high-quality and low-quality measurements as classified in Amidan et al. [2014]. (a) log ratio of quartile 2 to quartile 1 for MS1 spectra. (b) log ratio of quartile 2 to quartile 1 for MS2 spectra. (c) log ratio of quartile 3 to quartile 2 for MS1 spectra. (d) log ratio of quartile 3 to quartile 2 for MS2 spectra. (e) log ratio of quartile 4 to quartile 3 for MS1 spectra. (f) log ratio of quartile 4 to quartile 3 for MS2 spectra.

The Figures S3 and S4 demonstrate that the low-quality measurements (`poor`) have lower total ion current (TIC) values compared to high-quality measurements (`good`). The Figures serve as a visual check to differences in TIC values and aids in understanding the data quality of the measurements. It has to be pointed out that a further stratification (e.g. along the instrument type) might be helpful to further point out differences between the levels of data quality of the Amidan et al. [2014] data set.

The Figure S5 on the other hand does not indicate differences between data quality and might not be indicate of data quality for the quality issues of the Amidan et al. [2014] data set.

##### 3.3.1 Comparison to QuaMeter metrics

The XLS files also contain pre-calculated `QuaMeter` metrics [Ma et al., 2012] for each of the measurements. In the following, we will compare the `QuaMeter`-metrics to the `MsQuality` metrics to check if `MsQuality` shows concordant results compared to `QuaMeter`.

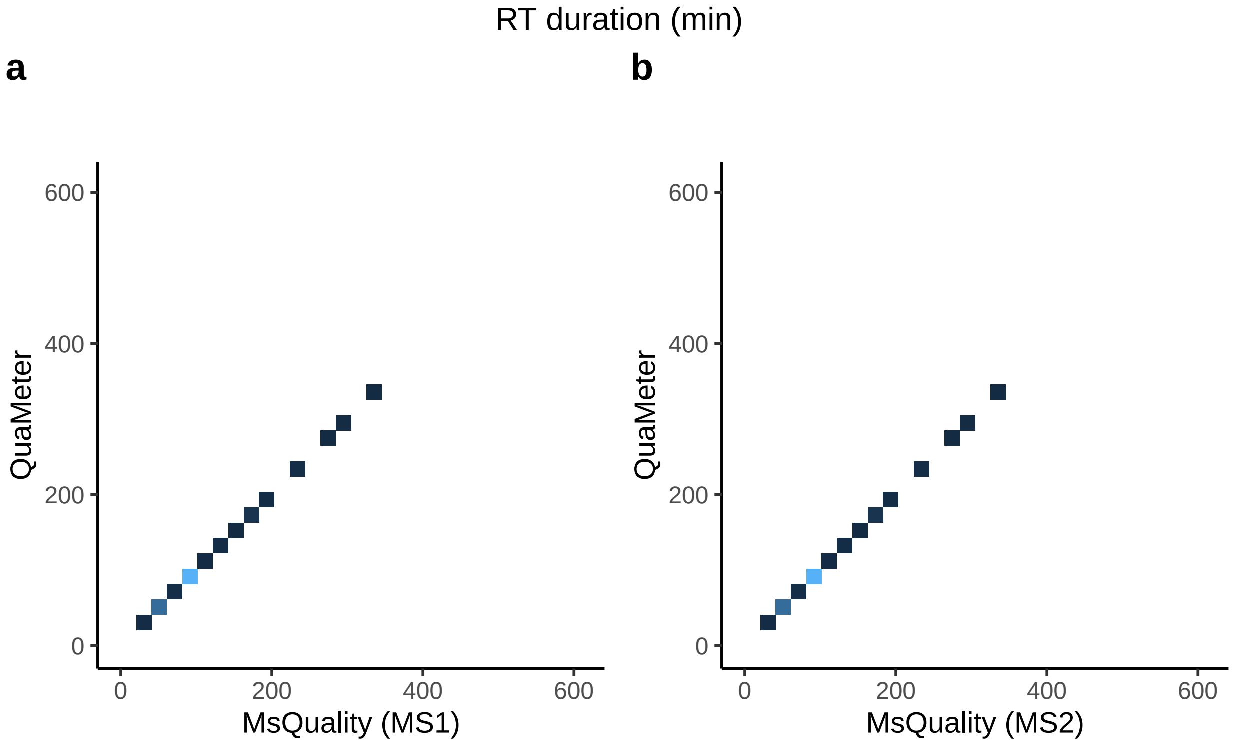

Figure S6: Comparison of quality metrics calculated by `MsQuality` and `QuaMeter`: RT duration (`rtDuration`). The corresponding metric for `QuaMeter` is `RT_Duration` (no specification if the metric was calculated on MS1 and/or MS2 spectra). The `MsQuality` metrics are calculated from MS1 and MS2 spectra. One data point is obtained per MS1 and MS2 spectra and the data points are displayed as 2D densities. Brighter areas correspond to high 2D density areas. (a) RT duration for MS1 spectra (`QuaMeter` metric: `RT_Duration`). (b) RT duration for MS2 spectra (`QuaMeter` metric: `RT_Duration`).

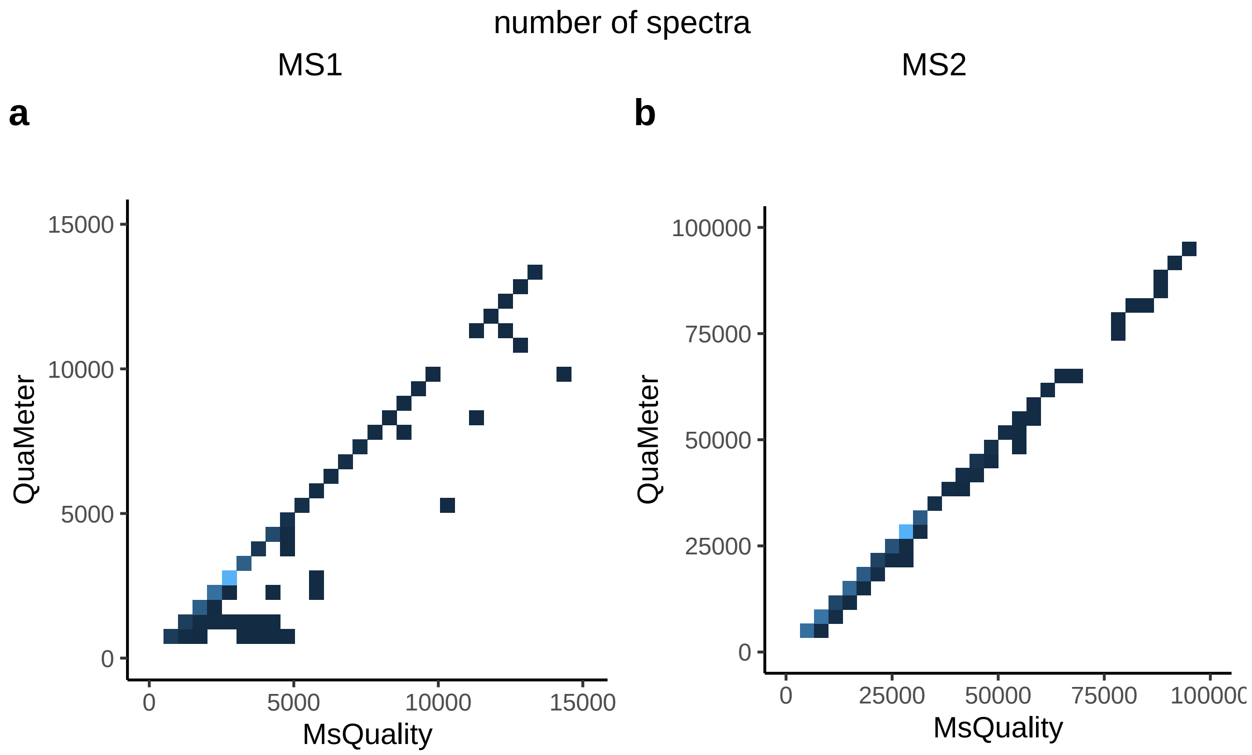

Figure S7: Comparison of quality metrics calculated by `MsQuality` and `QuaMeter`: Number of spectra (`numberSpectra`). The corresponding metrics for `QuaMeter` are `MS1_Count` and `MS2_Count`. The `MsQuality` metrics are calculated from MS1 and MS2 spectra. One data point is obtained per MS1 and MS2 spectra and the data points are displayed as 2D densities. Brighter areas correspond to high 2D density areas. (a) Number of MS1 spectra (`QuaMeter` metric: `MS1_Count`). (b) Number of MS2 spectra (`QuaMeter` metric: `MS2_Count`).

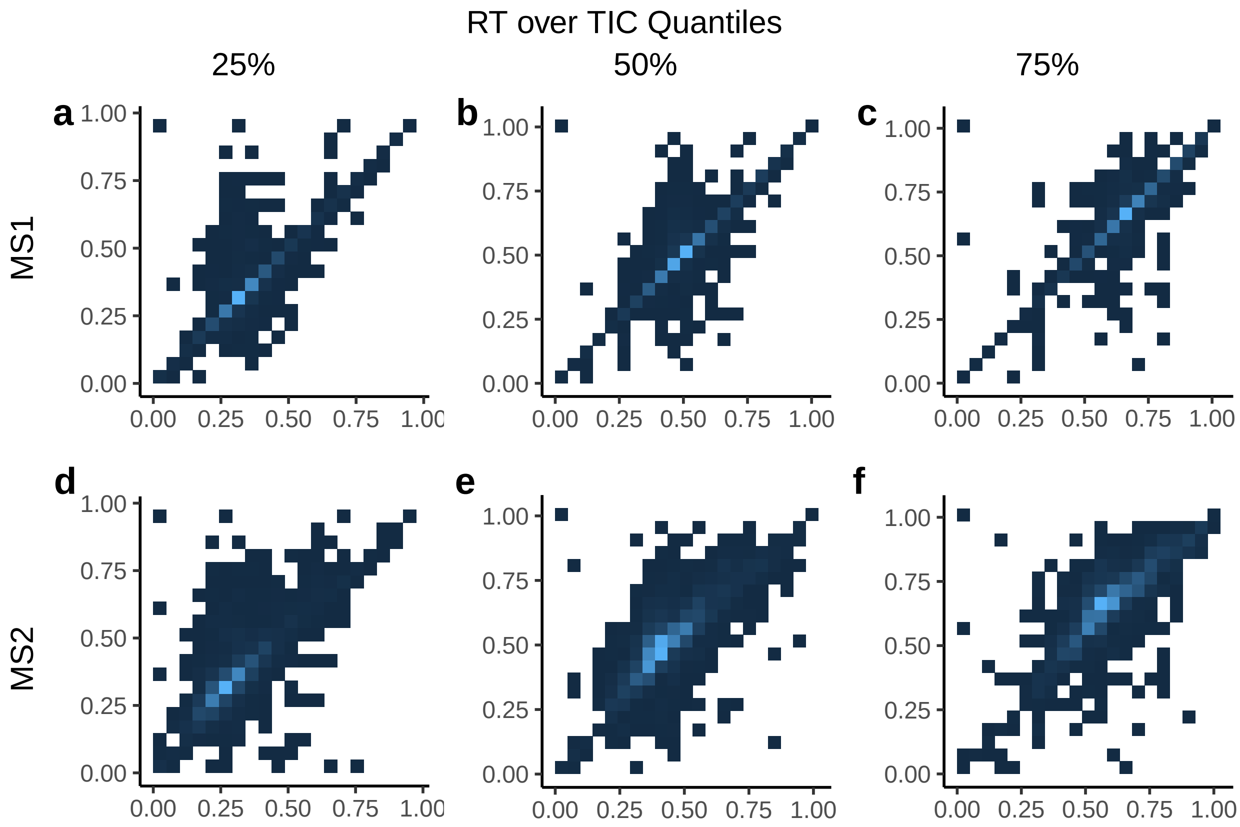

Figure S8: Scatterplot between quality metrics calculated by `MsQuality` and `QuaMeter`: RT over TIC quantile (`rtOverTicQuantiles`). The corresponding metrics for `QuaMeter` are `RT_TIC_Q1`, `RT_TIC_Q2`, and `RT_TIC_Q3` (no specification if these metrices were calculated on MS1 and/or MS2 spectra). The `MsQuality` metrics are calculated from MS1 and MS2 spectra. One data point is obtained per MS1 and MS2 spectra and the data points are displayed as 2D densities. Brighter areas correspond to high 2D density areas. (a) 25% quantile for MS1 spectra (`QuaMeter` metric: `RT_TIC_Q1`). (b) 50% quantile for MS1 spectra (`QuaMeter` metric: `RT_TIC_Q2`). (c) 75% quantile for MS1 spectra (`QuaMeter` metric: `RT_TIC_Q3`). (d) 25% quantile for MS2 spectra (`QuaMeter` metric: `RT_TIC_Q1`). (e) 50% quantile for MS2 spectra (`QuaMeter` metric: `RT_TIC_Q2`). (f) 75% quantile for MS2 spectra (`QuaMeter` metric: `RT_TIC_Q3`).

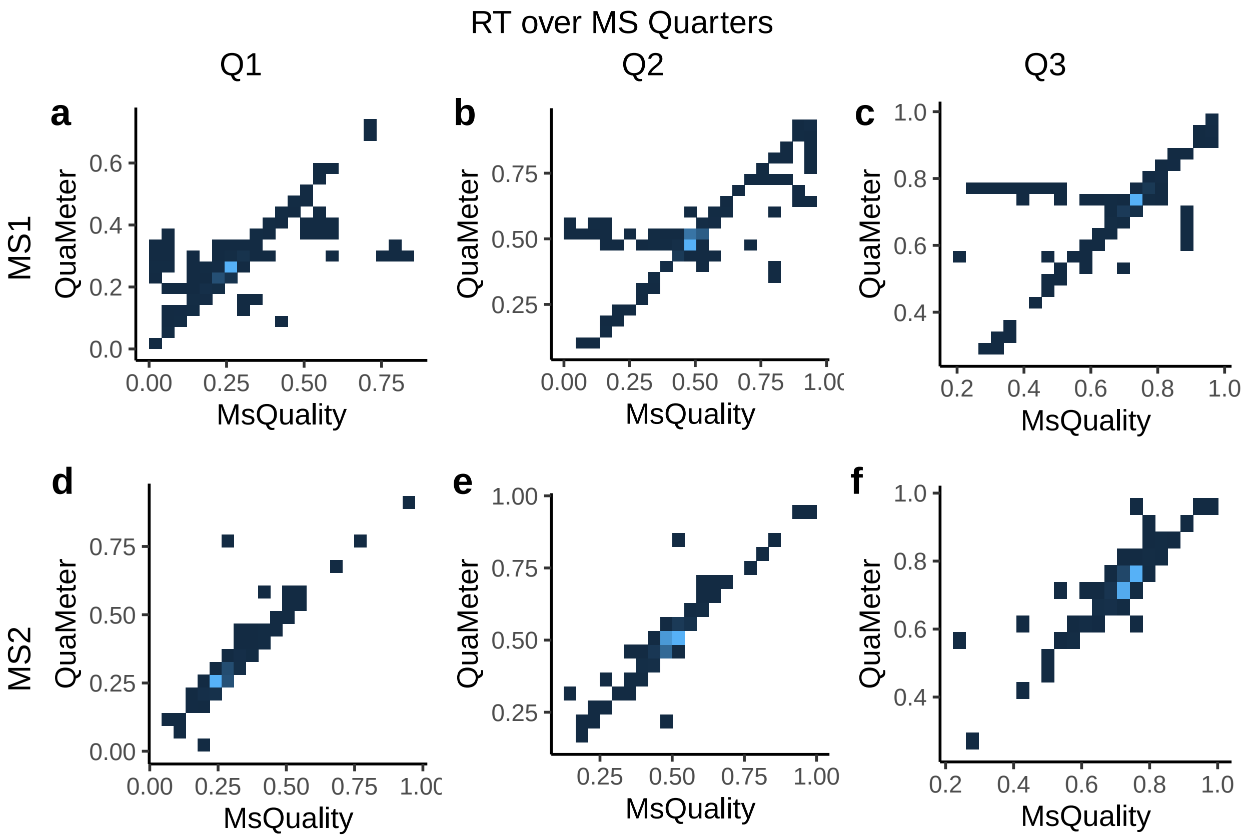

Figure S9: Comparison of quality metrics calculated by `MsQuality` and `QuaMeter`: RT over MS quarters (`rtOverMsQuarters`). The `MsQuality` metrics are calculated from MS1 and MS2 spectra. One data point is obtained per MS1 and MS2 spectra and the data points are displayed as 2D densities. Brighter areas correspond to high 2D density areas. (a) Quarter 1 for MS1 spectra. (b) Quarter 2 for MS1 spectra. (c) Quarter 3 for MS1 spectra. (d) Quarter 1 for MS2 spectra. (e) Quarter 2 for MS2 spectra. (f) Quarter 3 for MS2 spectra. Q: quarter.

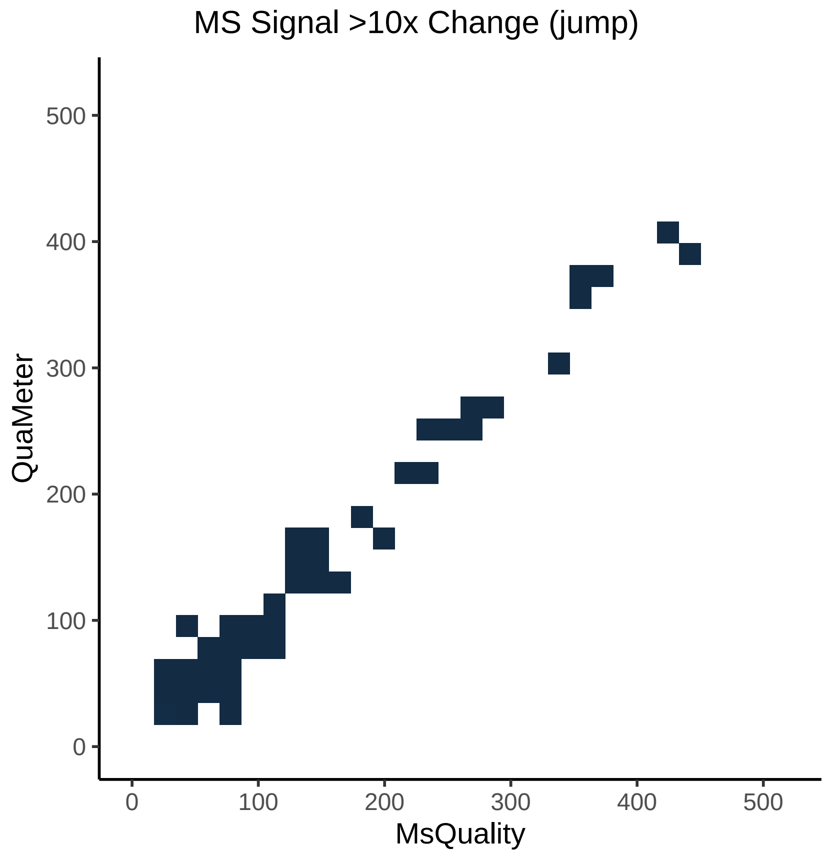

Figure S10: Comparison of quality metrics calculated by `MsQuality` and `QuaMeter`: MS Signal >10x Change (jump, `msSignal10xChange`). The metrics are only calculated from MS1 spectra. One data point is obtained per MS1 spectra and the data points are displayed as 2D densities. Brighter areas correspond to high 2D density areas.

The Figures S6, S7, S8, S9, S10 indicate that the `QuaMeter` and `MsQuality` generally compute similar values. This is shown by values that show high correlation (points locate close to the identity line within the scatter plots). We provide in Table S1 the Pearson and Spearman correlation coefficients of the `MsQuality` metrics with their corresponding `QuaMeter` metrics. Table S2 is a higher-level analysis regarding the quantiles of Pearson and Spearman correlation coefficients between the `MsQuality` and `QuaMeter` metrics. This analysis supports the observation that `MsQuality` calculates highly similar metric values as 75% of the metrics show Pearson/Spearman coefficients of 0.81/0.87 or higher.

Table S1: Pearson and Spearman correlation coefficients between `MsQuality` and pre-calculated QuaMeter metrics.

| MsQuality | QuaMeter | MS level | Pearson coef. | Spearman coef. |
|---|---|---|---|---|
| rtDuration | RT_Duration | 1 | 1 | 0.998 |
| numberSpectra | MS1_Count | 1 | 0.968 | 0.972 |
| numberSpectra | MS2_Count | 2 | 1 | 0.999 |
| rtOverTicQuantiles.25% | RT_TIC_Q1 | 1 | 0.887 | 0.884 |
| rtOverTicQuantiles.25% | RT_TIC_Q1 | 2 | 0.765 | 0.779 |
| rtOverTicQuantiles.50% | RT_TIC_Q2 | 1 | 0.902 | 0.888 |
| rtOverTicQuantiles.50% | RT_TIC_Q2 | 2 | 0.803 | 0.795 |
| rtOverTicQuantiles.75% | RT_TIC_Q3 | 1 | 0.909 | 0.912 |
| rtOverTicQuantiles.75% | RT_TIC_Q3 | 2 | 0.821 | 0.822 |
| rtOverMsQuarters.Quarter1 | RT_MS_Q1 | 1 | 0.734 | 0.908 |
| rtOverMsQuarters.Quarter1 | RT_MSMS_Q1 | 2 | 0.953 | 0.985 |
| rtOverMsQuarters.Quarter2 | RT_MS_Q2 | 1 | 0.806 | 0.9 |
| rtOverMsQuarters.Quarter2 | RT_MSMS_Q2 | 2 | 0.964 | 0.978 |
| rtOverMsQuarters.Quarter3 | RT_MS_Q3 | 1 | 0.812 | 0.916 |
| rtOverMsQuarters.Quarter3 | RT_MSMS_Q3 | 2 | 0.949 | 0.974 |
| msSignal10xChange | IS_1A | 1 | 0.687 | 0.838 |

Table S2: Quantiles for Pearson and Spearman correlation coefficients for `MsQuality` and pre-calculated QuaMeter metrics. The correlation analysis showed that 75% of the metrics showed Pearson/Spearman correlation coefficients over 0.81/0.87, 50% over 0.89/0.91, and 25% over 0.96/0.97 between `MsQuality` and QuaMeter metrics.

| Quantile | Pearson coef. | Spearman coef. |
|---|---:|---:|
| 0% | 0.69 | 0.78 |
| 10% | 0.75 | 0.81 |
| 20% | 0.80 | 0.84 |
| 25% | 0.81 | 0.87 |
| 30% | 0.81 | 0.89 |
| 40% | 0.82 | 0.90 |
| 50% | 0.89 | 0.91 |
| 60% | 0.91 | 0.92 |
| 70% | 0.95 | 0.97 |
| 75% | 0.96 | 0.97 |
| 80% | 0.96 | 0.98 |
| 90% | 0.98 | 0.99 |
| 100% | 1.00 | 1.00 |

#### 3.4 Performance under parallelization

Similar to the above-mentioned analysis using the flow injection analysis, an important aspect, especially when dealing with large amount of data, is scalability and performance when computing the quality metric.

We measure the time it takes to calculate the quality metrics under parallelization of the tasks on 1, 2, 4, 8, and 16 workers using the `microbenchmark` package. For computational reasons we limit the calculation to the first 500 .mzML files.

```
path <- "/scratch/naake/Amidan2014"
fls <- dir(path, full.names = TRUE, recursive = TRUE, pattern = "mzML") |>
    unique()
fls <- fls[1:500]
sps_mb <- sps[sps$dataOrigin %in% fls]

metrics <- c("rtDuration", "rtOverTicQuantiles", "rtOverMsQuarters",
    "ticQuartileToQuartileLogRatio", "numberSpectra",
    "medianPrecursorMz", "rtIqr", "rtIqrRate", "areaUnderTic",
    "areaUnderTicRtQuantiles", "medianTicRtIqr", "medianTicOfRtRange",
    "mzAcquisitionRange", "rtAcquisitionRange", "precursorIntensityRange",
    "precursorIntensityQuartiles", "precursorIntensityMean",
    "precursorIntensitySd", "msSignal10xChange", "ratioCharge1over2",
    "ratioCharge3over2", "ratioCharge4over2", "meanCharge",
    "medianCharge")

df_mb <- microbenchmark(
    workers_1 = calculateMetricsFromSpectra(spectra = sps_mb,
        metrics = metrics, BPPARAM = MulticoreParam(workers = 1)),
    workers_2 = calculateMetricsFromSpectra(spectra = sps_mb,
        metrics = metrics, BPPARAM = MulticoreParam(workers = 2)),
    workers_4 = calculateMetricsFromSpectra(spectra = sps_mb,
        metrics = metrics, BPPARAM = MulticoreParam(workers = 4)),
    workers_8 = calculateMetricsFromSpectra(spectra = sps_mb,
        metrics = metrics, BPPARAM = MulticoreParam(workers = 8)),
    workers_16 = calculateMetricsFromSpectra(spectra = sps_mb,
        metrics = metrics, BPPARAM = MulticoreParam(workers = 16)),
    times = 32L, control = list(warmup = 2), check = "equal"
)
```

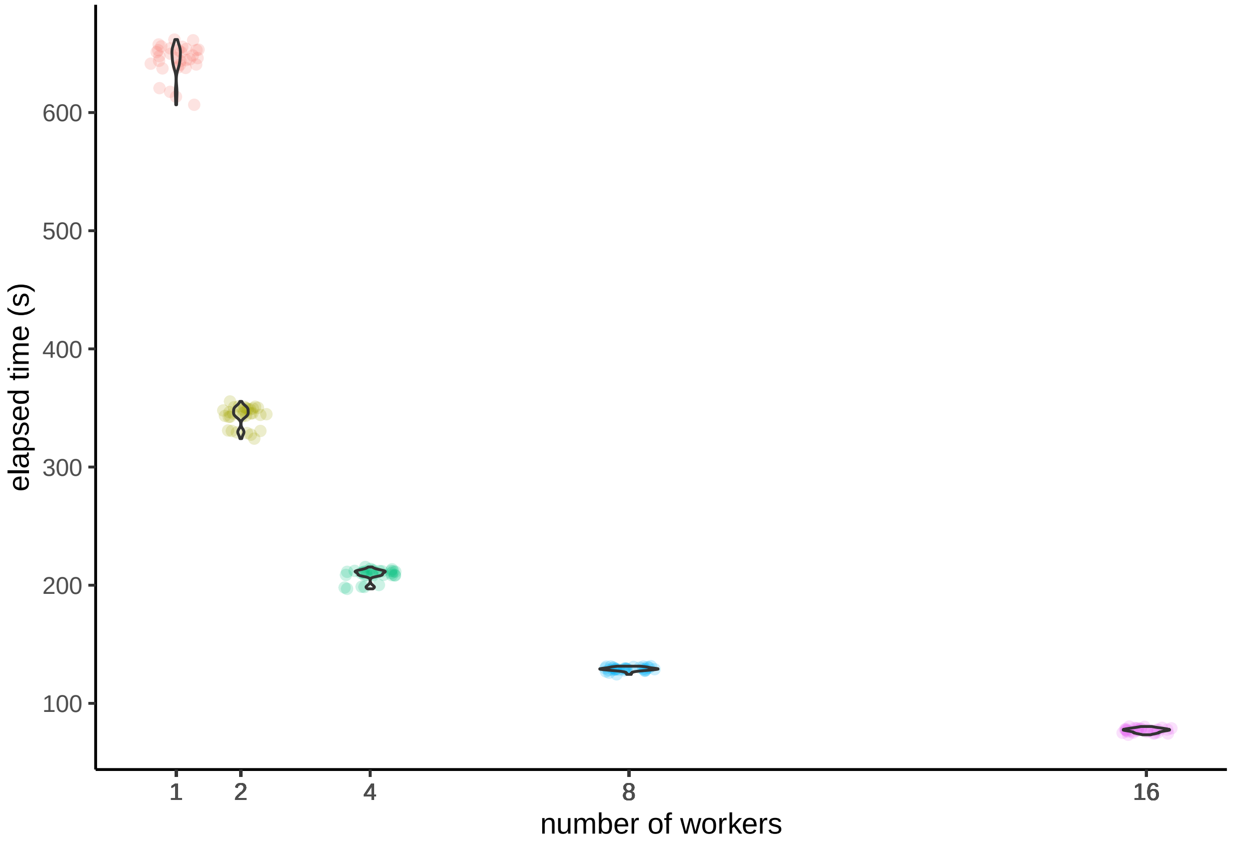

Figure S11: Execution time for the calculation of quality metrics of the data set of Amidan et al. [2014] under parallelization (1, 2, 4, 8, and 16 workers).

The `microbenchmark` package was used to accurately measure the performance improvements achieved by parallelization (Figure S11). By parallelizing the calculation of the quality metrics across multiple workers, it is possible to significantly reduce the execution time.

### 4 Session info

Information on the attached packages.

```
## R version 4.2.2 (2022-10-31 ucrt)
## Platform: x86_64-w64-mingw32/x64 (64-bit)
## Running under: Windows 10 x64 (build 19044)
##
## Matrix products: default
##
## locale:
## [1] LC_COLLATE=English_Germany.utf8  LC_CTYPE=English_Germany.utf8
## [3] LC_MONETARY=English_Germany.utf8 LC_NUMERIC=C
## [5] LC_TIME=English_Germany.utf8
##
## attached base packages:
## [1] stats4    stats     graphics  grDevices utils     datasets  methods
## [8] base
##
## other attached packages:
##  [1] patchwork_1.1.2      BiocFileCache_2.6.0  dbplyr_2.3.0
##  [4] curl_5.0.0           tidyr_1.3.0          tibble_3.1.8
##  [7] stringr_1.5.0        readxl_1.4.1         dplyr_1.1.0
## [10] ggpubr_0.5.0         ggbeeswarm_0.7.2     ggplot2_3.4.0
## [13] microbenchmark_1.4.9 MsQuality_0.99.6     mzR_2.32.0
## [16] Rcpp_1.0.10          Spectra_1.8.1        ProtGenerics_1.30.0
## [19] BiocParallel_1.32.5  S4Vectors_0.36.1     BiocGenerics_0.44.0
## [22] knitr_1.42           BiocStyle_2.26.0
```
